# Multiple resistance of *Colletotrichum truncatum* from soybean to QoI and MBC fungicides in Brazil

**DOI:** 10.1101/2022.06.24.497464

**Authors:** Flávia Rogério, Renata Rebellato Linhares de Castro, Nelson Sidnei Massola Júnior, Thaís Regina Boufleur, Ricardo Feliciano dos Santos

## Abstract

*Colletotrichum truncatum*, the most relevant fungal species associated with soybean anthracnose, is responsible for major losses in the crop. Chemical control via fungicide application is still the most effective strategy for the control of soybean foliar diseases. However, the increase in anthracnose incidence in some regions of Brazil indicates that current chemical control has not been effective against anthracnose. In this study, we assessed the fungicide sensitivity of *C. truncatum* lineages using isolates representing two important regions of soybean production in Brazil to the fungicides azoxystrobin, thiophanate-methyl, difenoconazole, and fludioxonil. We characterized the molecular resistance to quinone-outside inhibitors (QoI), methyl benzimidazole carbamates (MBC) and demethylation inhibitors (DMI) fungicide groups based on amino acid sequences of the cytochrome b (*cytb*), β-tubulin gene (β*-tub*), and P450 sterol 14a-demethylases (*CYP51*) genes. Multiple resistance of *C. truncatum* isolates to QoI and MBC was observed associated with mutation points in the β*-tub* (E198A and F200Y) and *cytb* (G143A). Alternatively, low EC_50_ values were found for fludioxonil and difenoconazole indicating high efficacy. Analysis of *C. truncatum* genomes revealed two potential DMI targets, *CYP51A* and *CYP51B*, and higher genetic variability in the *CYP51A* gene. A slight correlation between genetic differentiation of *C. truncatum* populations and fungicide sensibility was found (Student’s t-test <0.001). To our knowledge, this is the first report of multiple resistance to QoI and MBC fungicides in *C. truncatum* in Brazil.

**Highlights:**

- Multiple resistance of *C. truncatum* to azoxystrobin and thiophanate-methyl
- C. *truncatum* isolates are sensitive to difenoconazole and fludioxonil
- Presence of E198A and F200Y β*-tubulin* mutations and G143A *cytochrome b* mutation
- Presence of *CYP51A* and *CYP51B* paralogues and higher genetic variability in the *CYP51A*

## 1. Introduction

Soybean anthracnose caused by *Colletotrichum* species is one of the most important fungal diseases in the crop. Although new species have been reported recently associated with the disease, *Colletotrichum truncatum* is the most prominent species and is responsible for major losses in soybean fields (Boufleur et al., 2021; Shi et al., 2020). In Brazil, grain losses of 90 kg/ha of grain due to anthracnose were reported for each 1% increment in the incidence of the disease in commercial soybean fields (Dias et al., 2016). However, the disease can lead to total crop loss under favorable weather conditions of high temperature and moisture (EMBRAPA, 2008; Yang and Hartman, 2016).

Since the emergence of Asian soybean rust (*Phakopsora pachyrhizi*), anthracnose has been underestimated in Brazil. Frequent reports on the increase of soybean anthracnose in the North and Central-West regions indicate that chemical control program for fungal diseases in soybean, focusing mainly on rust, has not been effective against anthracnose (Dias et al., 2016). Considering that most of the soybean production in Brazil derives from the aforementioned regions (CONAB, 2020), which present optimal weather conditions for disease development, losses by anthracnose threaten national production.

Chemical control (including seed treatment and fungicide application) is still the most effective strategy for anthracnose management. A large number of commercial products belonging to different chemical groups are registered for soybean anthracnose control in the country (AGROFIT, 2021); however, little information on their efficacy is available. Most commercial products employed for soybean fungal diseases control are mixtures of single active ingredients, which belong in the majority to the chemical group quinone-outside inhibitors (QoI), methyl benzimidazole carbamates (MBC), demethylation inhibitors (DMI), phenylpyrrole (PP), and succinate dehydrogenase inhibitors (SDHI) (FRAC, 2021; Pesqueira et al., 2016).

The increase in the use of fungicides, especially through repetitive applications of molecules with the single-site mode of action, can imply a high selection pressure for resistance. Some studies reported losses of sensitivity of *C. truncatum* isolates to QoI, MBC, and DMI from different crops (Chen et al., 2018, 2016; Dias et al., 2016; Poti et al., 2020; Torres-Calzada et al., 2015). In Brazil studies indicate that anthracnose chemical control has not been satisfactory in soybean fields in Tocantins State, with maximum efficiency of only 41.7% for azoxystrobin (QoI) and cyproconazole (DMI), suggesting that other regions with a similar microclimate of humidity and temperature may be at risk (Dias et al., 2016, 2019).

QoI, MBC, and DMI are widely used in agriculture and have specific modes of action, in contrast to multi-site inhibitors that act in a wide range of cellular processes (FRAC, 2021). Site-specific fungicides are conducive to resistance selection since a single mutation on the target protein can cause resistance, and thus loss of effectiveness (Ma and Michailides, 2005). Although fungicide resistance can be conferred by different mechanisms, the majority is due to substitutions in amino acid sequences of the target proteins (Ma and Michailides, 2005; Mair et al., 2016). Molecular investigation of resistance on target sites responsible for fungicide efficiency is a useful approach to detect resistant fungal genotypes and allows optimize their management (Lucas et al., 2015).

QoI fungicides inhibit fungal mitochondrial respiration by binding the ubiquinol-oxidizing (Qo) site of cytochrome *b (cytb)*, therefore blocking electrons transport and preventing ATP production (Bartlett et al., 2002). Three amino acid substitutions are associated with QoI resistance (F129L, G137R, and G143A), responsible for different levels of resistance (Gisi et al., 2002; Lucas et al., 2015). MBC fungicides act by inhibiting cell division binding to the beta-tubulin (β*-tub*) gene, and preventing microtubule assembly, disrupting chromosome segregation and migration (Brennan et al., 2007; Downing, 2000). Several target site mutations are associated with resistance, mostly in the codons E198A/G/K and F200Y (FRAC, 2021).

DMI resistance involves the disruption of fungal growth by inhibition of the gene cytochrome P450 sterol 14α-demethylase (*CYP51*) in the biosynthesis of sterol (Ziogas and Malandrakis, 2015). The mechanisms of resistance to this group are poorly understood, but three processes underlying resistance have been documented: (i) target-site modification in the gene *CYP51* (Délye et al., 1998), *CYP51* overexpression (Hamamoto et al., 2000), and increased drug efflux pumps (Sanglard et al., 1995). Mutations in the *CYP51* gene seem to be the major mechanism (Cools et al., 2013), and several pathogenic fungi such as *Uncinula necator, Blumeria graminis, Erysiphe graminis*, and *Candida albicans* exhibited mutation in resistant isolates (Délye et al., 1998, 1997; Favre et al., 1999; Wyand and Brown, 2005). *Colletotrichum* species can possess two paralogous *CYP51* genes, which showed different levels of sensitivity to DMI fungicides (Chen et al., 2020; Wang et al., 2020; Wei et al., 2020). Fludioxonil, a phenylpyrrole (PP) fungicide, has a speculative mechanism of action (FRAC 2021). Although resistance to the group is classified as low to medium risk, resistance was identified in other fungal species (Iacomi-Vasilescu et al., 2004; Kanetis et al., 2008). Previous studies demonstrated fludioxonil efficacy against *C. acutatum* (Wedge et al., 2007), but no information about *C. truncatum* is available.

Fungal plant pathogens frequently show genetically divergent lineages as a consequence of populational subdivision, caused by distinct factors, like geographic distance and host specialization (James et al., 2006; Soanes et al., 2007). Different lineages may have different mechanisms to cause diseases gained during the co-evolutionary arms race between fungal populations and their hosts (Plissonneau et al., 2017; Van Oosterhout, 2021). In other words, distinct populations may hold different virulence factors. *C. truncatum* is thought to be an invasive species introduced in Brazil multiple times, which led to the establishment of three genetic lineages spread throughout soybean fields (Rogério et al., 2022, 2019). These lineages possess different levels of genetic variation with evidence of sexual recombination, which could bolster levels of adaptation to soybean cultivation.

Due to the increase of anthracnose importance in Brazil, this study aimed to investigate the sensitivity of *C. truncatum* isolates from important soybean production regions to four fungicides (azoxystrobin, thiophanate-methyl, difenoconazole, and fludioxonil) and perform molecular characterization of isolates with different levels of sensibility to these fungicides.

## 2. Material and Methods

### 2.1. Fungal isolates

Isolates used in this study were collected in 2016 and 2017, from ten soybean commercial fields in two Brazilian regions showing a high incidence of anthracnose (Table 1). These isolates were previously genotyped by multilocus microsatellite typing and whole-genome sequencing (Rogério et al., 2022, 2019), and they are representative of the three genetic groups (C1, C2, and C3) detected in those fields.

**Table 1.** *Colletotrichum truncatum* isolates used in this study.

| Lineage | Isolate | State | GenBank accession number |  |  |  |
| --- | --- | --- | --- | --- | --- | --- |
|  |  |  | *TUB2* | *CYTB* | *CYP51A* | *CYP51B* |
| C1 | LFN0169 | Mato Grosso | MZ682550 | MZ682567 | MZ682584 | MZ682601 |
| C1 | LFN0185 | Mato Grosso | MZ682551 | MZ682568 | MZ682585 | MZ682602 |
| C1 | LFN0262 | Mato Grosso | MZ682556 | MZ682573 | MZ682590 | MZ682607 |
| C1 | LFN0309 | Goiás | MZ682544 | MZ682561 | MZ682578 | MZ682595 |
| C1 | LFN0360 | Goiás | MZ682548 | MZ682565 | MZ682582 | MZ682599 |
| C1 | LFN0297 | Goiás | MZ682542 | MZ682559 | MZ682576 | MZ682593 |
| C1 | LFN0346 | Goiás | MZ682546 | MZ682563 | MZ682580 | MZ682597 |
| C2 | LFN0205 | Mato Grosso | MZ682552 | MZ682569 | MZ682586 | MZ682603 |
| C2 | LFN0217 | Mato Grosso | MZ682553 | MZ682570 | MZ682587 | MZ682604 |
| C2 | LFN0248 | Mato Grosso | MZ682555 | MZ682572 | MZ682589 | MZ682606 |
| C2 | LFN0318 | Goiás | MZ682545 | MZ682562 | MZ682579 | MZ682596 |
| C2 | LFN0349 | Goiás | MZ682547 | MZ682564 | MZ682581 | MZ682598 |
| C2 | LFN0288 | Goiás | MZ682541 | MZ682558 | MZ682575 | MZ682592 |
| C3 | LFN0150 | Mato Grosso | MZ682549 | MZ682566 | MZ682583 | MZ682600 |
| C3 | LFN0225 | Mato Grosso | MZ682554 | MZ682571 | MZ682588 | MZ682605 |
| C3 | LFN0268 | Mato Grosso | MZ682557 | MZ682574 | MZ682591 | MZ682608 |
| C3 | LFN0308 | Goiás | MZ682543 | MZ682560 | MZ682577 | MZ682594 |

### 2.2. In vitro fungicide sensitivity assays

The sensitivity of *C. truncatum* isolates to fungicides was determined based on mycelial growth inhibition assay fungicide-amended on potato dextrose agar medium (PDA). We used commercial formulations of azoxystrobin (Amistar 500 WG, Syngenta Crop Protection), thiophanate-methyl (Cercobin 700 WP, Ihara), difenoconazole (Score 250 EC, Syngenta Crop Protection), and fludioxonil (Maxim 25, Syngenta Crop Protection). These fungicides were selected based on the active ingredients registered for soybean anthracnose control in Brazil (AGROFIT, 2021).

Based on preliminary assays, we observed that *C. truncatum* isolates showed intense mycelial growth, with an effective concentration to inhibit 50% of the mycelial growth (EC_50_) higher than 100 μg/ml for azoxystrobin and thiophanate-methyl fungicides. In this way, we used single discriminatory doses of 100 μg/ml to distinguish between resistant and sensitive isolates for these fungicides. Isolates that showed a percentage of mycelial growth inhibition higher than 50% were classified as resistant. Five-millimeter-diameter mycelial plugs were taken from actively growing 7-day-old colonies on PDA and transferred to PDA plates amended with the fungicide concentration of 0 and 100 μg/ml. Plates were incubated at 25°C under constant light for 5 days. Each fungicide-isolate combination and control plate (i.e., plates onto non-amended PDA) were replicated three times and experiments were performed twice. The diameter of each colony was used to calculate the percentage of mycelium inhibition (MGI). MGI was obtained using the formula: MGI = ((C-FT) / C) * 100), where MGI is the mycelial growth inhibition, C is the control treatment colony diameter and FT is the fungicide treatment colony diameter.

The sensitivity of *C. truncatum* isolates to the difenoconazole and fludioxonil was also determined by mycelia growth assays. Mycelia plugs were placed upside down onto PDA dishes amended with difenoconazole at 0, 0.01, 0.1, and 10 µg active ingredient (a.i.)/ml; and Fludioxonil at 0, 0.001, 0.01, 0.1, and 1 µg active ingredient (a.i.)/ml. The experiment was performed following the methodology described above. A regression analysis based on the percentage of mycelial growth inhibition was performed to estimate the EC_50_ value for these fungicides. The experiment was performed twice, and the combined data demonstrated that variances were homogeneous according to *F-test* (*P* < 0.05).

### 2.3. Molecular characterization of fungicides target genes

To investigate point mutations in the *cytb*, β*-tub*, and *CYP51* genes we used genomic data available from all isolates (Rogério et al., 2022). The BLASTn tool (Altschul et al., 1990) was used to retrieve the gene sequences related to resistance to the fungicides from the genomes. The *cytb* and β*-tub* genes were retrieved from genomes using as query sequences the strain *C. truncatum* CMES1059 (GenBank accession number MK163913.1 and MK188497, respectively). For the DMI group, the presence of the paralogs *CYP51A* and *CYP51B* was investigated, as well as the point mutations on them. Therefore, we used the strain *C. truncatum* CtRR131 as query sequences (GenBank accession number MG799553.1 and MG799552, respectively). Predicted amino acid sequences along the DNA sequences obtained were performed using Expasy Bioinformatics Resource Portal and aligned using MEGA11 software (Kumar et al., 2016).

### 2.4. Phylogenetic analysis

The deduced amino acid sequences of *CYP51* paralogs genes were used to investigate the phylogenetic relationship between isolates. A phylogenetic tree was constructed based on the concatenated alignment of *CYP51* sequences of *C. truncatum* generated in this study, in addition to *CYP51* homologs from several *Colletotrichum* species and other closely related ascomycete fungi, including *Saccharomyces cerevisiae* as outgroup (GenBank accession XP003713527.1). Multiple alignments were performed using MAFFT v. 7.490 (Katoh et al., 2002) implemented in Geneious 8.1.4., (http://www.geneious.com). The phylogenetic analysis was conducted by the maximum likelihood (ML) method using the JTT matrix-based model.

## 3. Results

### 3.1. Fungicide sensitivity *in vitro* assays

The isolates were tested using a single dose of 100 μg/ml of azoxystrobin and thiophanate-methyl, which differentiated resistant from sensitive isolates to both fungicides. On the first hand, all isolates were resistant to azoxystrobin (growth inhibition up to 22%) while for thiophanate-methyl only the isolates LFN0217 (lineage C2) and LFN0225 (Lineage C3) were sensitive, showing a percentage of mycelial growth inhibition 65 and 88%. On the other hand, all isolates were sensitive to difenoconazole and fludioxonil, with EC_50_ values ranging from 0.06 to 0.61 μg mL^-1^ (mean of 0.17 μg mL^-1^) to difenoconazole, and 0.21 e 2.97 μg mL^-1^ (mean of 0.84 μg mL^-1^) to fludioxonil (Table 2).

**Table 2.** Sensitivity of *Colletotrichum truncatum* isolates from soybean in Brazil to fludioxonil and difenoconazole fungicides.

| Mean EC<sub>50</sub> values (µg/ml) |  |  |  |
| --- | --- | --- | --- |
| Lineage | Isolate | Fludioxonil | Difenoconazole |
| C1 | LFN0297 | 0.078 | 1.244 |
| C1 | LFN0346 | 0.211 | 1.422 |
| C1 | LFN0360 | 0.178 | 1.523 |
| C1 | LFN0309 | 0.081 | 1.035 |
| C1 | LFN0169 | 0.089 | 1.150 |
| C1 | LFN0185 | 0.464 | 2.968 |
| C1 | LFN0262 | 0.081 | 0.526 |
| C2 | LFN0318 | 0.079 | 1.245 |
| C2 | LFN0217 | 0.145 | 0.440 |
| C2 | LFN0248 | 0.055 | 0.233 |
| C2 | LFN0205 | 0.083 | 0.221 |
| C2 | LFN0288 | 0.088 | 0.668 |
| C2 | LFN0349 | 0.182 | 0.312 |
| C3 | LFN0150 | 0.607 | 0.227 |
| C3 | LFN0308 | 0.053 | 0.650 |
| C3 | LFN0268 | 0.131 | 0.212 |
| C3 | LFN0225 | 0.371 | 0.283 |

### 3.2. Molecular characterization of fungicide resistance mutations

The nucleotide sequences translated from the *cytb* gene of the 17 *C. truncatum* isolates revealed a substitution from glycine (G) to alanine (A) at codon 143 in all isolates analyzed (Fig.1). This mutation is well documented in the literature to confers resistance to QoI fungicides. These isolates were classified as resistant based on *in vitro* sensibility assay and such resistance was supported at the molecular level.

**Figure 1.**
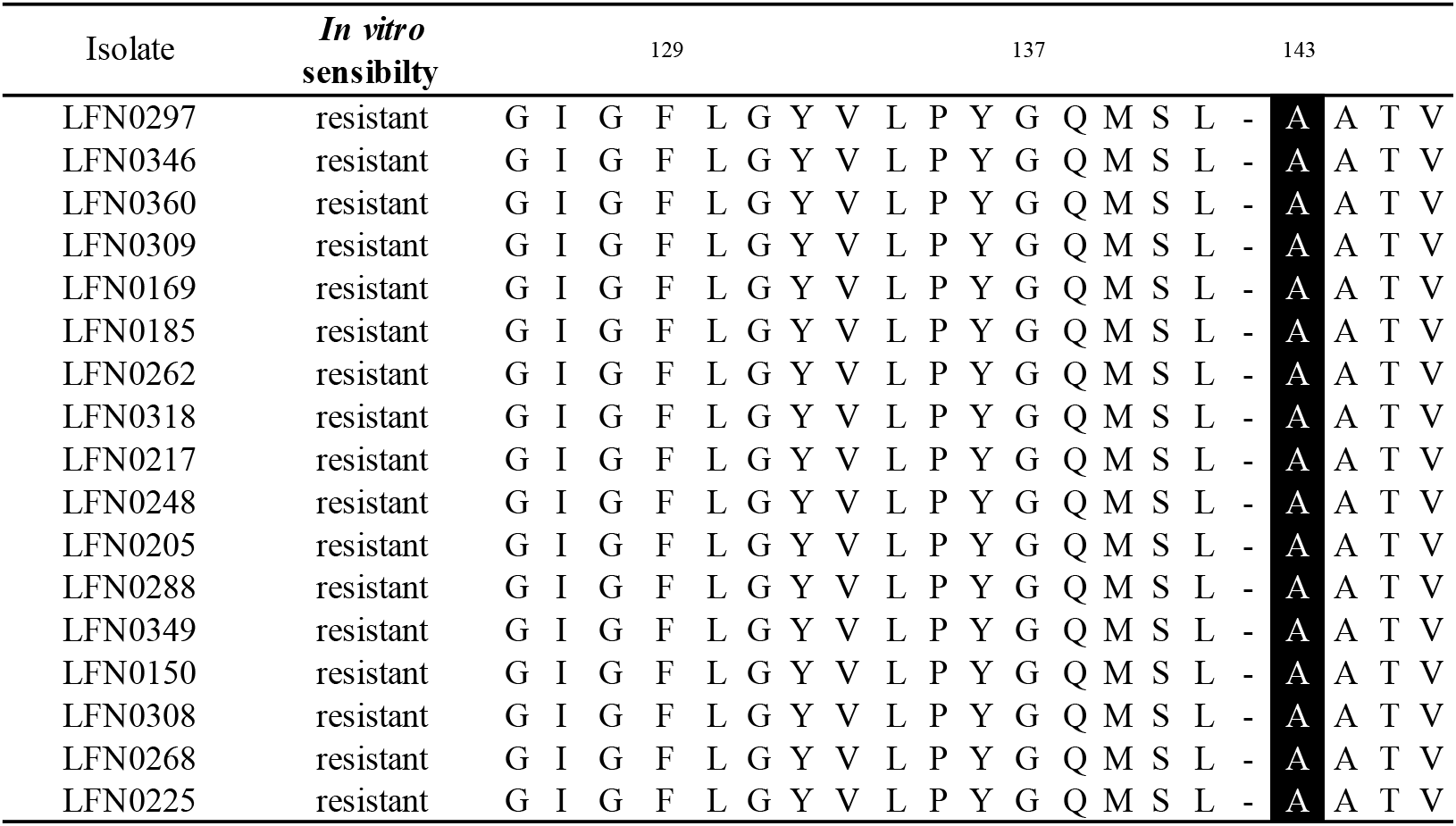
Aligned amino acid sequences of partial cytochrome b gene (codons 126 to 146) of *Colletotrichum truncatum* isolates from soybean. The mutation associated with QoI resistance was observed at codon 143. Amino acids: A - alanine; F - phenylalanine; G - glycine; I - isoleucine; L - leucine; M - methionine; N - asparagine; P - proline; Q - glutamine; S - serine; T - threonine; V - valine; W - tryptophan; Y - tyrosine.

Analysis of β*-tub* gene sequences revealed mutations at codons 198 and 200, which confers resistance to MBC fungicides, confirming the sensitivity obtained *in vitro* tests. The isolates LFN0346, LFN360, LFN390, LFN169, LFN185, LFN318, LFN248, LFN205, LFN349, LFN150, and LFN308 showed substitutions from glutamic (E) to alanine (A) at codon 198 (E198A), while the isolates LFN0297, LFN262, LFN288, and LFN268 showed substitutions from phenylalanine (F) to tyrosine (Y) at codon 200 (F200Y). In contrast, the isolates LFN217 (lineage C2) and LFN225 (lineage C3), which were sensitive to thiophanate-methyl in the *in vitro* tests, did not exhibit any of these mutations (Fig.2).

**Figure 2.**
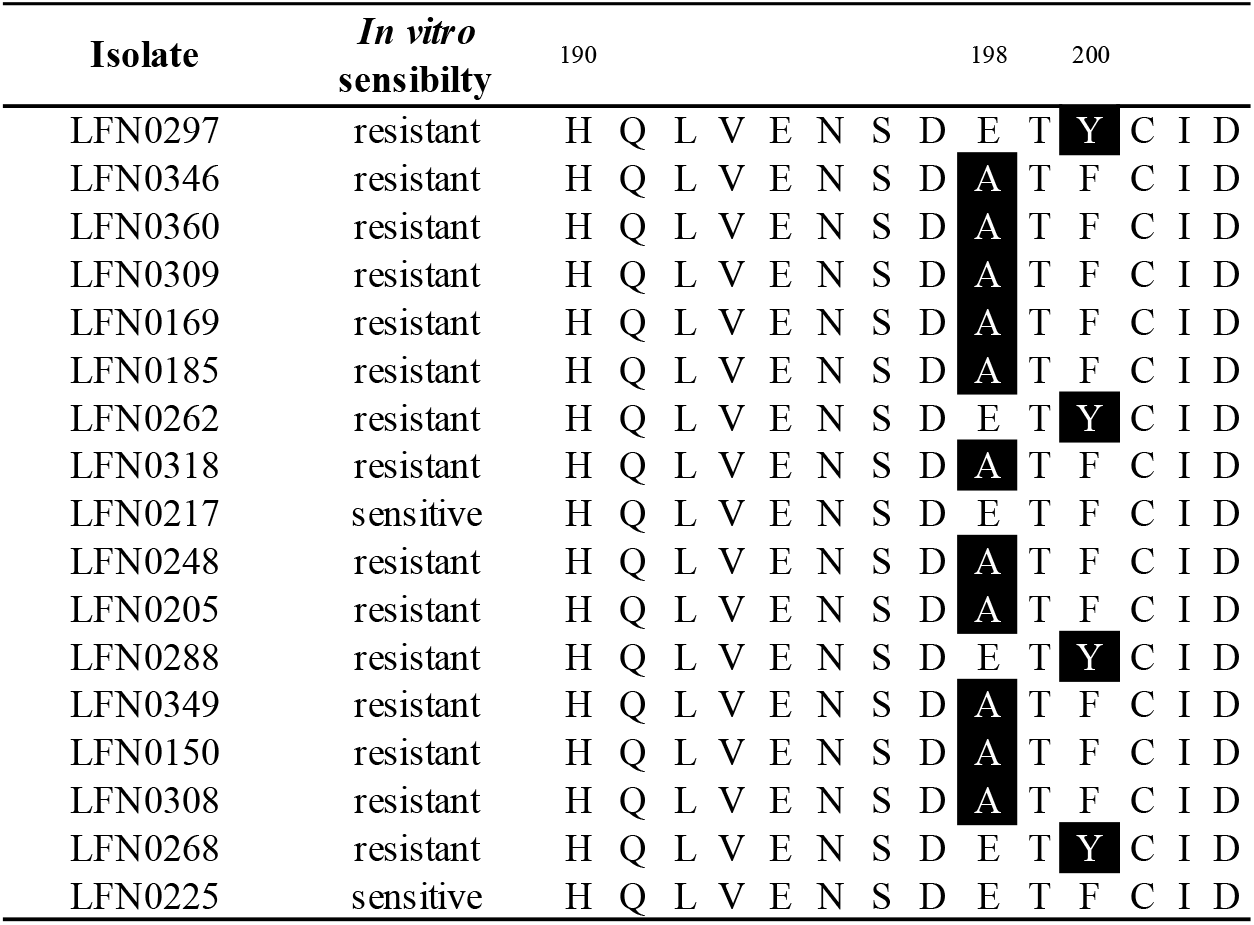
Aligned amino acid sequences of partial beta-tubulin gene (codons 190 to 200) of *Colletotrichum truncatum* from soybean. Highlighted in black are mutations at codon E198A and F200 linked to MBC resistance. Amino acids: A - alanine; D - acid aspartic; E - acid glutamic; F - phenylalanine; H - histidine; I - isoleucine; L - leucine; N - asparagine; P - proline; Q - glutamine; S - serine; T - threonine; V - valine.

### 3.3. Phylogenetic inference of *CYP51* gene

Analysis of *C. truncatum* genomes to DMI resistance revealed two paralogous *CYP51* genes, *CYP51A* and *CYP51B*, that putatively encode the protein P450 sterol 14a-demethylases (Fig.3). The deduced 512 amino-acid protein encoded by the 1,539 bp coding sequence from *CYP51A* and the 526 amino-acid protein encoded by the 1,578 bp coding sequence from *CYP51B* were analyzed regarding the presence of mutations.

The substrate recognition sites (SRS) in *CYP51* genes are very conserved in filamentous fungi, and the amino acid alterations occurring around the azole-binding site of the enzyme affect its affinity, and they are commonly investigated in DMI resistance (Han et al., 2010; Mellado et al., 2001). The alignment of sequences from *C. truncatum* isolates and *Aspergillus fumigatus*, here used as reference (GenBank accession number XP_752137.1), revealed eight variations in amino acid sequences, present in 3 SRS, in the form of E105D (SRS1), D253Q, D280E (SRS4), L391V, K484C, K484S, P501A(SRS6) and P501T (SRS6) for *CYP51A* gene (Fig. S1). For *CYP51B* no variations were detected between isolates.

**Figure 3.**
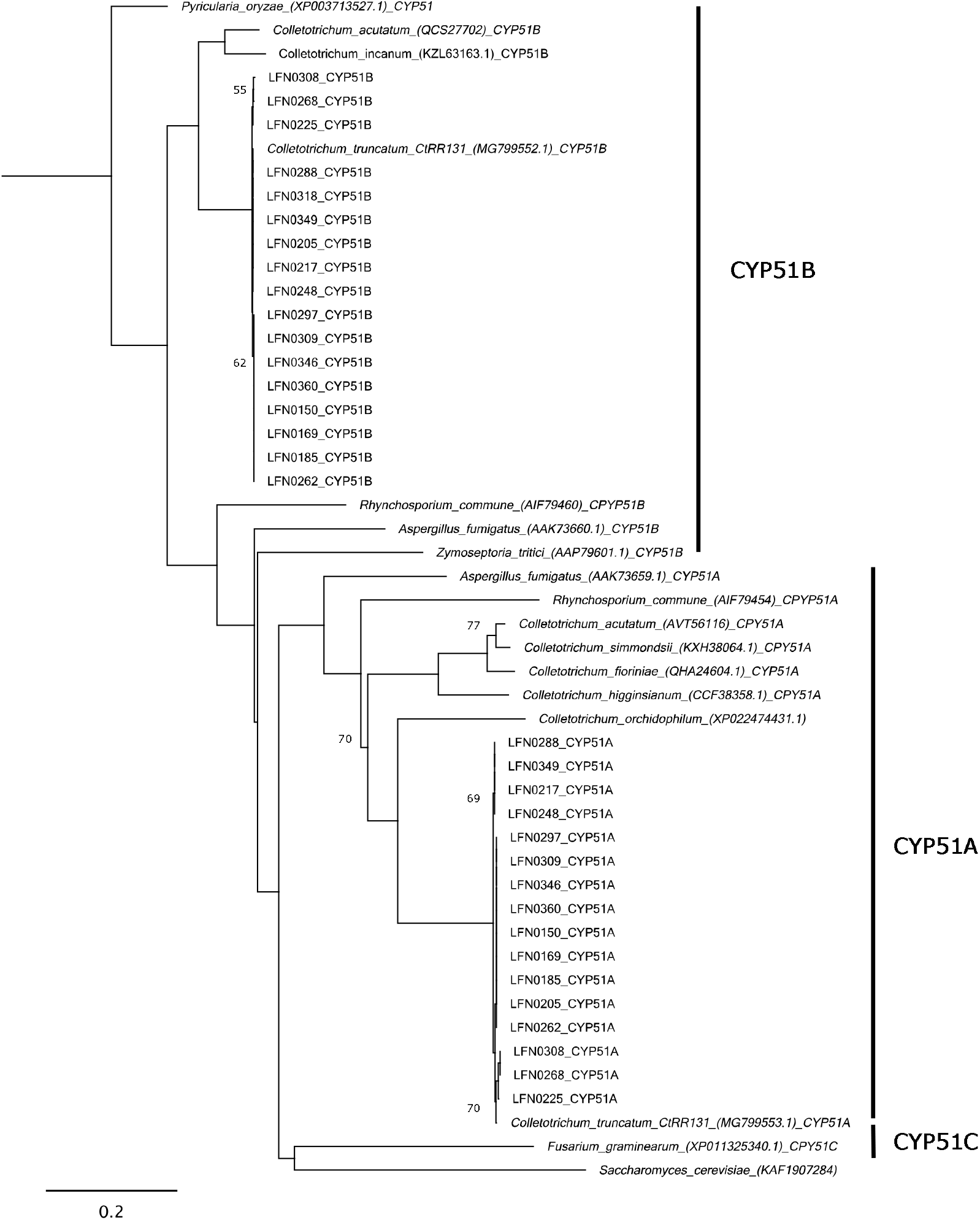
Phylogenetic inference of *CYP51* proteins generated by the maximum likelihood method. The deduced amino acid sequences of seventeen *Colletotrichum truncatum* isolates from soybean and other *Colletotrichum* species and fungal species were used in this analysis: <u>CYP51A</u> - *C. truncatum* (strain CtRR131) (GenBank accession nº MG799553.1); *C. acutatum* (GenBank accession nº AVT56116); *C. simmondsii* (GenBank accession nº KXH38064.1); *C. higginsianum* (GenBank accession nº CCF38358.1); *C. orchidophilum* (GenBank accession nº XP022474431.1); *C. fioriniae* (GenBank accession nº QHA24604.1); *Rhynchosporium commune* (GenBank accession nº AIF79454). <u>CYP51B</u>-*Aspergillus fumigatus* (GenBank accession nº AAK73659.1); *C. truncatum* (strain CtRR131) (GenBank accession nº MG799552.1*); C. acutatum* (GenBank accession nº QCS27702); *C. incanum* (GenBank accession nº KZL63163.1); *Rhynchosporium commune* (GenBank accession nº AIF79460); *Aspergillus fumigatus* (GenBank accession nº AAK73660.1); *Zymoseptoria tritici* (GenBank accession nº AAP79601.1); <u>CYP51C</u>-*Fusarium graminearum* (GenBank accession nº XP011325340.1); CYP51-*Pyricularia oryzae* (GenBank accession nº XP003713527.1); *Saccharomyces cerevisiae* (GenBank accession nº KAF1907284). Support values below 80 are shown on the nodes.

## 4. Discussion

The recent increase in soybean anthracnose importance in some regions of Brazil indicates that current chemical management employed for fungal disease control has not been effective against anthracnose. Although chemical control is the main method of anthracnose control, little information about its effectiveness is available. Here, we reported *in vitro* resistance of *C. truncatum* isolates to azoxystrobin and thiophanate-methyl associated with point mutations in the *cytb* (G143A) and β*-tub* (E198A and F200Y) genes. Multiple resistance to QoI and MBC were also recently reported in *Corynespora cassiicola*, another important soybean fungus in Brazil, showing the same mutations detected in this study (Mello et al., 2022). Alternatively, low EC_50_ values were found for fludioxonil and difenoconazole indicating high efficacy. We detected the presence of two *CYP51* paralogous (*CYP51A* and *CYP51B*) and higher genetic variability in the *CYP51A* gene. A slight correlation between genetic differentiation of *C. truncatum* populations and fungicide sensibility was found. Difenoconazole EC_50_ values for lineage C1 were statistically different from other lineages (Student’s t-test <0.001).

All isolates showed *in vitro* resistance to azoxystrobin (QoI) and thiophanate-methyl (MBC) fungicides using a single discriminatory dose of 100 μg/ml. Such phenotypic responses were supported at the molecular level. Analysis of the *cytb* gene revealed presence of the G143A mutation in all isolates. QoI resistance has been detected in several *Colletotrichum* species such as *C. graminicola, C. siamense, C. acutatum*, and *C. cereale* (Avila-Adame et al., 2003; Chechi et al., 2019; Forcelini et al., 2016; Hu et al., 2015; Young et al., 2010). For *C. truncatum*, isolates highly resistant to azoxystrobin were already reported, but the molecular mechanism conferring resistance was not investigated (Torres-Calzada et al., 2015). To our knowledge, this study is the first report of G143A mutation associated with QoI resistance in this species.

For thiophanate-methyl, we found both resistant and sensitive isolates, being resistance predominant (88%). All resistant isolates showed F200Y mutations in the β*-tub* gene. Mutation at codon 198 is mostly found in isolates with high levels of MBC resistance, while a mutation at position 200 is correlated with moderate levels (Lucas et al., 2015). Resistance to MBC fungicides has already been reported in *C. truncatum* from different crops. A high frequency of isolates resistant to carbendazim was observed in soybean fields in Thailand, with the presence of both mutations (Poti et al., 2020). Isolates from several hosts (including pepper, papaya, and physic nut) also showed resistance to thiabendazole associated with the E198A mutation (Torres-Calzada et al., 2015). A previous study investigated the efficacy of several fungicides (including carbendazim) to soybean anthracnose under natural conditions in Brazil, and they concluded that fungicide efficacy is gradually being reduced against anthracnose (Dias et al., 2016). The high risk of MBC resistance in *Colletotrichum* spp. is recognized and should be considered to anthracnose control (Nalumpang et al., 2010; Suwan and Na-Lampang, 2013; Torres-Calzada et al., 2015; Vieira et al., 2017; Wong et al., 2008). Cross-resistance between MBC fungicides is reported in several phytopathogenic fungi (Chung et al., 2010; Cunha and Rizzo, 2003; Sun et al., 2010; Wong et al., 2008), and it can represent a risk to chemical control since MBCs have been widely used in soybean fields for a long time, either alone or in mixture with other fungicide groups such as DMIs and QoIs (Pesqueira et al., 2016).

Regarding sensibility for difenoconazole and fludioxonil, belonging to DMI and SDHI, all isolates were sensitive to both fungicides. The low EC_50_ value is consistent with previous studies with difenoconazole in *C. truncatum* (Chen et al., 2018, 2016; Zhang et al., 2017). We detected the presence of two *CYP51* paralogous, but no known mutations associated with DMI resistance were revealed. A similar result was also found in *C. gloeosporioides* isolates evaluated to difenoconazole and propiconazole (Wang et al., 2020). However, we detected eight variations in amino acid sequences in 3 substrate recognition sites (SRS) in the *CYP51A* gene. In contrast, no variations were detected in *CYP51B*, in concordance with previous studies which say that variation at this paralogous is unusual (Brunner et al., 2015; Délye et al., 1997). The *CYP51A* gene is reported as more relevant to DMI sensitivity, and its higher variability suggests that it can adapt more rapidly under selection pressure, suggesting positive diversifying selection, while *CYP51B* takes over the more conserved function, and purifying selection can be acting on it (Brunner et al., 2015; Chen et al., 2018). Some studies have reported resistance of *C. truncatum* from several hosts (including soybean) to DMI fungicides (Carstens et al., 2017; Chen et al., 2018, 2016; Zhang et al., 2017). These findings point to the inherent resistance of *C. truncatum* to DMI fungicides. Although in this study we did not find resistance, the presence of high nucleotide variability in the *CYP51A* gene may indicate ongoing selective pressure on it, suggesting a risk to the development of resistance.

According to our results, the genetic differentiation of *C. truncatum* populations had not impacted the fungicide sensibility to the fungicides investigated. We expected that lineage C3, which is largely affected by genetic introgression from other lineages, comprising a high amount of secreted protein-encoding genes obtained by such genetic exchanges may exhibit higher virulence factors, for instance, fungicide mutation conferring resistance (Rogério et al., 2022). In other words, the genetic structure of the fungus present in the soybean fields is not related to different phenotypic fungicide sensibility. However, we found a significant difference (Student’s t-test <0.001) in EC_50_ values to difenoconazole for the lineage C1. Correlation among population structure and phenotypes is the most challenging to evolutionary genetics studies because virulence factors are associated with single genes, and subdivisional populational is a genome-wide effect. In conclusion, our study reveals multiple resistance of *C. truncatum* to QoI and DMI fungicides, widely used in soybean fields, for the first time in Brazil.

## CRediT authorship contribution statement

**Flávia Rogério:** Conceived the study, designed the project, performed analyzes, and wrote the manuscript. **Renata Rebellato Linhares de Castro:** Performed analyses and reviewed the manuscript.**Nelson Sidnei Massola Júnior:** Conceived the study, designed the project, and reviewed the manuscript. **Thaís Regina Boufleur:** Performed analyses and reviewed the manuscript. **Ricardo Feliciano dos Santos:** Performed analyses and reviewed the manuscript. All authors have read and agreed to the published version of the manuscript.

## Acknowledgments

This work was supported by the São Paulo Research Foundation (FAPESP 2017/09178-8), National Science and Technology Development Council (CNPq 305289/2018-7), and National Council for the Improvement of Higher Education (PROEX/CAPES 330002037002P3). Flávia Rogério received a Postdoctoral Research Fellowship from CAPES (Higher Education Personnel Improvement Coordination, Brazil) from “Estágio Pós-Doutoral do Programa Nacional de Pós-Doutorado” (PNPD - 2019-1293).

