## Supplementary material for "Multiple resistance of *Colletotrichum truncatum* from soybean to QoI and MBC fungicides in Brazil": Figure S1

Figure S1. Alignments of *CYP51A* amino acid sequences showing substrate recognition sites (SRS) from *Aspergillus fumigatus* (used as reference) and *Colletotrichum truncatum* isolates from soybean.

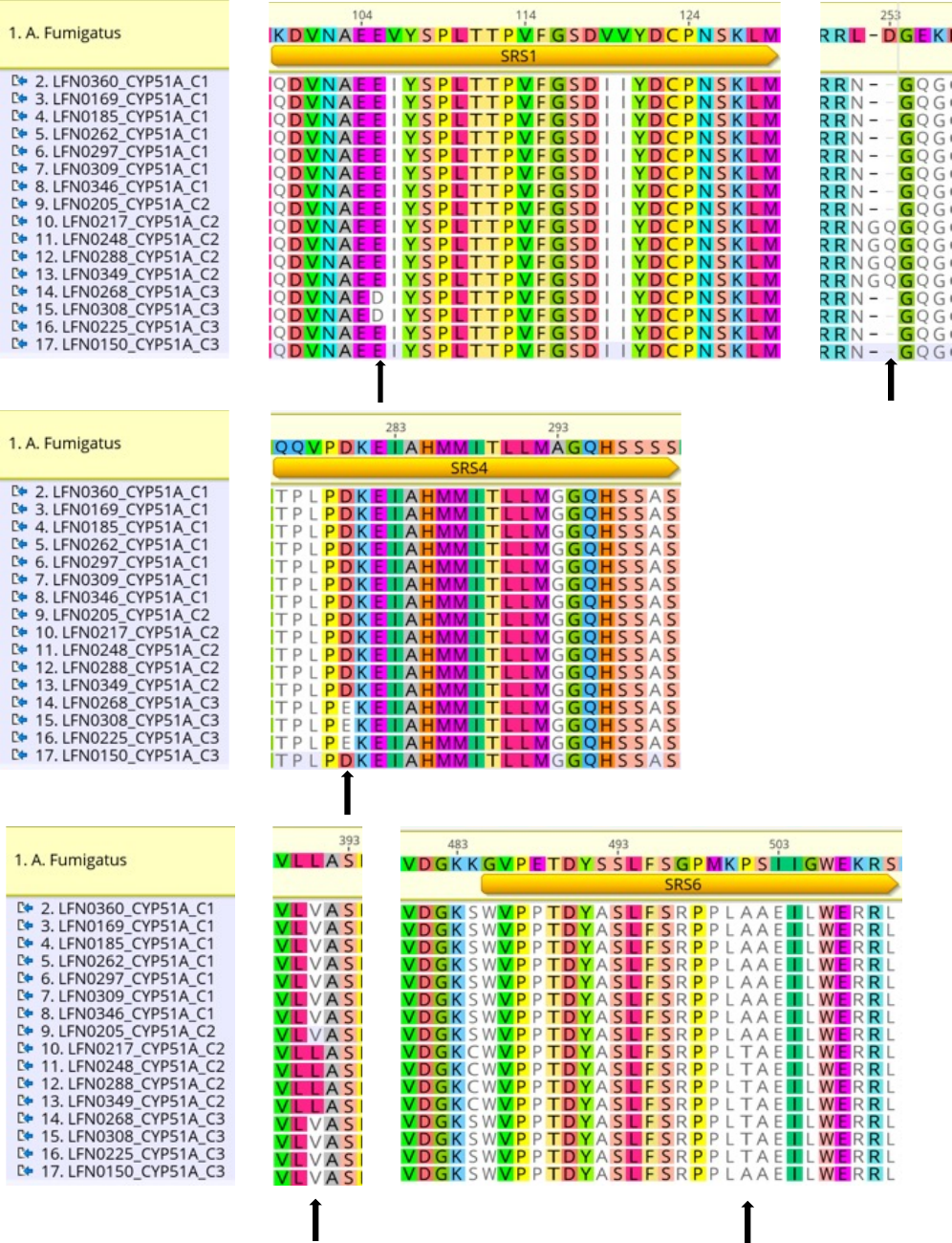
